# Multigenerational exposure to elevated temperatures leads to a reduction in standard metabolic rate in the wild

**DOI:** 10.1101/749986

**Authors:** Natalie Pilakouta, Shaun S. Killen, Bjarni K. Kristjánsson, Skúli Skúlason, Jan Lindström, Neil B. Metcalfe, Kevin J. Parsons

## Abstract

1. In light of global climate change, there is a pressing need to understand and predict the capacity of populations to respond to rising temperatures. Metabolic rate is a key trait that is likely to influence the ability to cope with climate change. Yet, empirical and theoretical work on metabolic rate responses to temperature changes has so far produced mixed results and conflicting predictions.
2. Our study addresses this issue using a novel approach of comparing fish populations in geothermally warmed lakes and adjacent ambient-temperature lakes in Iceland. This unique ‘natural experiment’ provides repeated and independent examples of populations experiencing contrasting thermal environments for many generations over a small geographic scale, thereby avoiding the confounding factors associated with latitudinal or elevational comparisons. Using Icelandic sticklebacks from three warm and three cold habitats, we measured individual metabolic rates across a range of acclimation temperatures to obtain reaction norms for each population.
3. We found a general pattern for a lower standard metabolic rate in sticklebacks from warm habitats when measured at a common temperature, as predicted by Krogh’s rule. Metabolic rate differences between warm- and cold-habitat sticklebacks were more pronounced at more extreme acclimation temperatures, suggesting the release of cryptic genetic variation upon exposure to novel conditions, which can reveal hidden evolutionary potential. We also found a stronger divergence in metabolic rate between thermal habitats in allopatry than sympatry, indicating that gene flow may constrain physiological adaptation when dispersal between warm and cold habitats is possible.
4. In sum, our study suggests that fish may diverge toward a lower standard metabolic rate in a warming world, but this might depend on connectivity and gene flow between different thermal habitats.

## Introduction

Climate change poses a substantial threat to biodiversity, as rising temperatures are altering biotic and abiotic environmental conditions and imposing novel selection pressures on organisms (Crozier & Hutchings 2014). In light of this, there is a pressing need to understand the capacity of populations to respond and adapt to increasing temperatures. This knowledge is critical for our ability to predict the impacts of climate change and to inform management and conservation efforts (Donelson et al. 2011, Sinclair et al. 2016, Campbell et al. 2017). Ectothermic animals are particularly vulnerable to changes in ambient temperature, because this directly influences their body temperature (Zuo et al. 2012). Ectotherms are therefore expected to adapt to climate change through plastic and/or evolutionary changes in a wide range of physiological, morphological, and behavioural traits (Crozier & Hutchings 2014).

A key trait that is likely to influence the capacity of ectothermic animals to cope with increasing temperatures is metabolic rate (Donelson et al. 2012). Standard and maximum metabolic rates are among the most commonly measured physiological traits and can have important fitness implications for organisms (White and Kearney 2013, Pettersen et al. 2018). Standard metabolic rate (SMR) refers to the minimum rate of energy throughput needed to sustain an animal at rest, whereas maximum metabolic rate (MMR) sets the upper limit for the capacity to perform oxygen-consuming physiological activities (Killen et al. 2016). Another important physiological variable is absolute aerobic scope (AS), which is calculated as the difference between an individual’s SMR and MMR. AS represents an individual’s total capacity for simultaneous oxygen-consuming tasks above maintenance requirements (e.g., digestion, activity, growth; Pörtner & Farrell 2008). In ectotherms, AS may increase with temperature up to an optimum, after which it declines with further temperature increases. AS has therefore been suggested to play a key role in the response of fish populations to changing temperatures (Farrell 2016, Sandblom et al. 2016). Notably, however, many fish species appear to show little or no change in AS with changes in temperature (Lefevre 2016, Nati et al. 2016, Jutfelt et al. 2018), so the degree to which AS limits adaptive responses to warming remains an open question.

It is also still unclear how SMR and MMR will evolve in response to climate change. According to Krogh’s rule (also known as metabolic cold adaptation), ectotherms living in cold environments should have higher metabolic rates than those in warm environments, when the two are observed at the same temperature (Krogh 1916, Gaston et al. 2009). This view is in line with the concept of countergradient variation, which is a result of stabilising selection favouring similar phenotypes in different environments (Marcil et al. 2006) and can arise when genetic differences in a trait counteract environmental effects (Conover & Schultz 1995). An alternative hypothesis is that natural selection will favour a reduction in metabolic rate in animals living at low temperatures if the energetic cost of cellular respiration exceeds the benefit of additional ATP production (Clarke 1980, Clarke 1991, Clark 1993, Chown and Gaston 1999). These opposing predictions have led to much controversy and debate about the effects of temperature changes on metabolic rate adaptation (Huey and Berrigan 1996, Chown and Gaston 1999, Addo-Bediako et al. 2002).

Until now, empirical studies aiming to address this issue have focused on interspecific and intraspecific variation in metabolic rate across contrasting thermal habitats, such as populations from different latitudes or elevations (e.g., Tsuji 1988, White et al. 2012, Gaitán-Espitia and Nespolo 2014; but see Sandblom et al. 2016). However, there are a number of confounding factors associated with these comparisons. For example, high-latitude habitats are not only colder on average but also experience more extreme temporal fluctuations in temperature and photoperiod than low-latitude habitats. These confounding factors may partly explain the mixed results of previous research on the effects of thermal environment on metabolic rate in ectotherms (Clarke 2003, White and Kearney 2013, Alton et al. 2017). For instance, high-latitude *Drosophila melanogaster* populations tend to have elevated metabolic rates compared to low-latitude populations as predicted by Krogh’s rule (Berrigan & Partridge 1997), but latitudinal comparisons of snail populations and altitudinal comparisons of isopod populations offer no support for this rule (Lardies et al. 2004, Gaitán-Espitia and Nespolo 2014). Another approach that has been used to study metabolic rate adaptation is experimental evolution. One such study found that *D. melanogaster* lines maintained at a low temperature evolved an elevated metabolic rate, consistent with Krogh’s rule (Berrigan & Partridge 1997). Nevertheless, a recent study on *D. melanogaster* found no evidence that cold environments select for a higher metabolic rate (Alton et al. 2017). Large-scale comparative studies examining the relationship between thermal environment and metabolic rate have also produced mixed results. Global-scale analyses of 346 insect species (Addo-Bediako et al. 2002) and 37 fish species (White et al. 2011) found support for Krogh’s rule, but an analysis of 65 Drosophilidae species did not (Messamah et al. 2017).

Here, we address this question using a novel approach of comparing populations of threespine sticklebacks (*Gasterosteus aculeatus*) found in geothermally warmed lakes and adjacent ambient-temperature lakes in Iceland. This unique ‘natural experiment’ provides repeated and independent examples of populations experiencing contrasting thermal environments for many generations over a small geographic scale, thereby avoiding the confounding factors associated with latitudinal or elevational comparisons. In addition, there are no substantial differences in water chemistry between geothermally warmed and ambient-temperature lakes, allowing us to isolate the effects of temperature. We have previously shown that there is strong morphological divergence between sticklebacks from these warm and cold habitats, suggesting local adaptation (Pilakouta et al. 2019 preprint).

Using sticklebacks from three warm and three cold habitats, we measured individual metabolic rates across a range of acclimation temperatures (10°C, 15°C, and 20°C) to obtain reaction norms for each population. We examined whether living in a warm environment over several generations results in a suppressed metabolic rate, providing a powerful test of the controversial Krogh’s rule. If metabolic rate responds in a similar way to thermal habitat across multiple populations, this would suggest that metabolic rate evolution in response to climate change may be predictable (Bolnick et al. 2018). We also investigated whether the steepness of metabolic rate reaction norms differs between populations from warm vs cold environments. Lastly, measuring metabolic rate at unfamiliar temperatures (i.e., 10°C for warm-habitat fish and 20°C for cold-habitat fish) allowed us to detect the release of cryptic genetic variation and thus hidden evolutionary potential to respond to thermal changes (Paaby & Rockman 2014, Shama 2017). Given the potential importance of physiological adaptation for population persistence (Donelson et al. 2011, Sinclair et al. 2016), our study could provide valuable insights into the capacity of ectotherms to cope with global climate change.

## Methods

### Study populations

Using unbaited minnow traps, we collected adult threespine sticklebacks from six freshwater populations in Iceland in May–June 2016 (Table 1). Two of these populations were allopatric, meaning that the warm and cold habitats were in neighbouring but separate water bodies with no potential for gene flow (Figure 1). Because these populations were located in close proximity to the marine habitat (and were thus more likely to be directly invaded by a common marine ancestor), we considered them a comparable warm-cold ‘population pair’. We also sampled two sympatric warm-cold population pairs, where the warm and cold habitats were in the same water body, with no physical barriers between them (Figure 1). The cold habitats have all existed for thousands of years, since the last glacial period (Einarsson et al. 2004), but there is some variation in the age of the warm habitats (Table 1). The ‘Mývatn warm’ and Grettislaug sites have been naturally heated by geothermal activity for over 2000 years (Hight 1965, Einarsson 1982). In contrast, the ‘Áshildarholtsvatn warm’ habitat originated only 50–70 years ago, fed by excess hot water runoff from nearby residences using geothermal heating. Since the generation time for threespine sticklebacks is about 1 year, the age of the warm habitats corresponds to the maximum number of generations each population pair may have been separated (Table 1).

**Table 1.** Sampling locations of warm- and cold-habitat threespine sticklebacks collected in May–June 2016 in Iceland. Distance refers to how far apart the warm-habitat and cold-habitat sample sites are for each warm-cold pair. All cold habitats have existed since the last glacial period and are therefore approximately 10,000 years old, whereas warm habitats can be classified as either young (<70 years old) or old (>2,000 years old). The summer and winter temperatures listed are the average water temperatures recorded at each sampling location during the corresponding seasons.

| Population pair | Water body | Thermal habitat | Age of warm habitat (years) | Distance (km) | Summer temperature (°C) | Winter temperature (°C) |
| --- | --- | --- | --- | --- | --- | --- |
| allopatric populations | Grettislaug | warm | >2,000 | 21.04 | 24.9 | 13.5 |
|  | Garðsvatn | cold |  |  | 14.6 | 2.2 |
| sympatric population 1 | Áshildarholtsvatn | warm | 50–70 | 0.05 | 24.1 | 12.5 |
|  |  | cold |  |  | 12.2 | 3.4 |
| sympatric population 2 | Mývatn | warm | >2,000 | 3.18 | 22.8 | 22.0 |
|  |  | cold |  |  | 11.5 | 1.0 |

**Figure 1.**
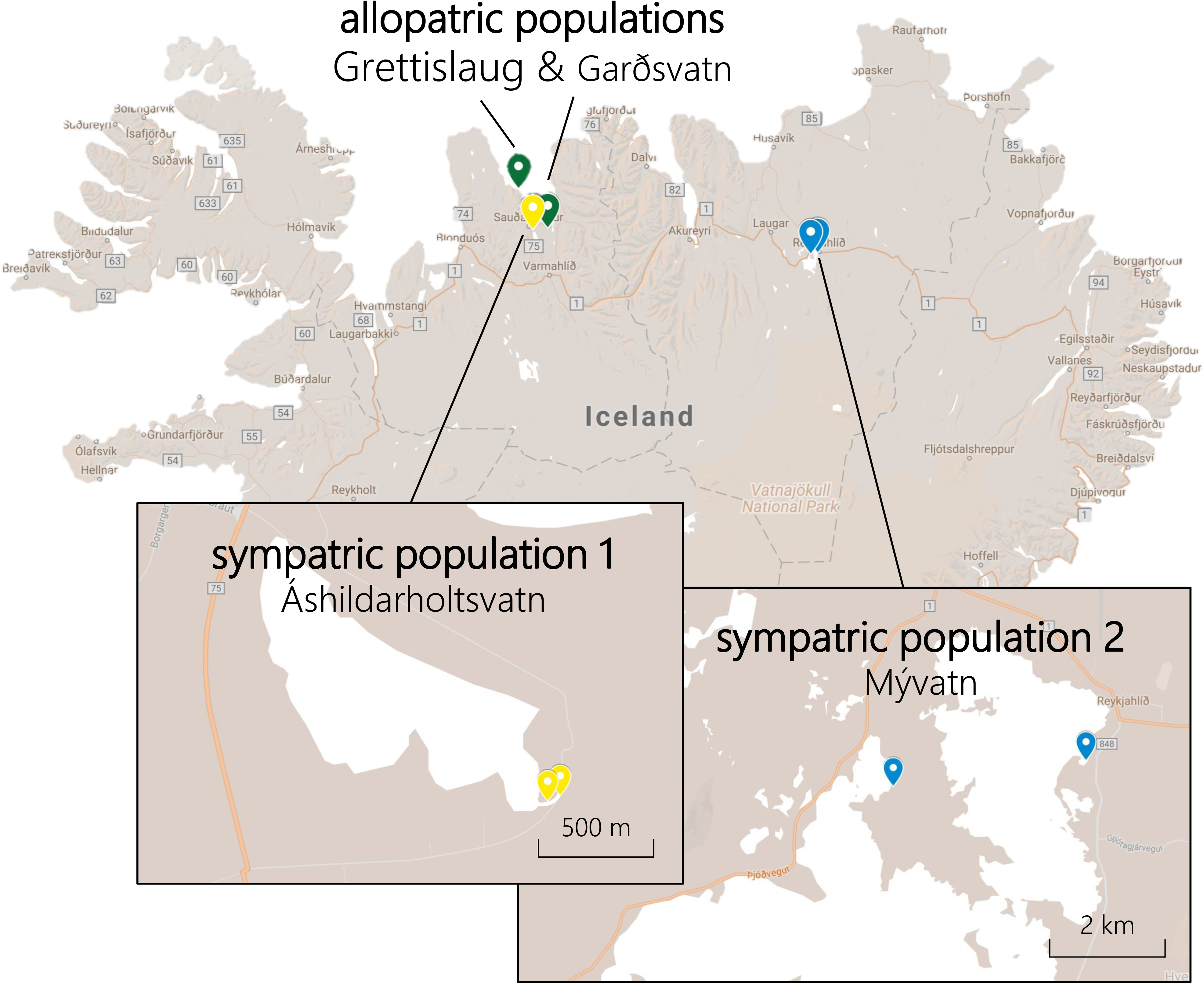
Map of Iceland showing the sampling locations of warm and cold habitat sticklebacks we collected for this study. Each of the three population pairs is indicated by a different colour.

### Transport of study animals

Before transport to University of Glasgow, we fasted sticklebacks for 48 hr to minimise the build up of ammonia in the transport water. On the day of shipping, we placed approximately 100 sticklebacks in each 100-litre polyethylene bag containing 25 litres of water. Air was removed from the bags and replaced with pure oxygen. Bags were sealed and placed inside insulated Styrofoam shipping boxes to minimise temperature fluctuations during transport. The fish were in transit for approximately 72 hr before arriving in Glasgow. No mortality was observed during transport.

### Animal husbandry

Once these fish arrived at University of Glasgow, they were kept at densities of 10-15 individuals per 10-litre tank in a common recirculation system at 15°C. The tanks contained plastic plants as shelter. Fish were fed *ad libitum* twice a day with a mixture of frozen bloodworms, *Mysis* shrimp, and *Daphnia*. Two months before the experiments, all fish were anaesthetised using benzocaine and marked with visible implant elastomer tags (Northwest Marine Technology Inc) to allow individual identification. They were kept at a 12h light:12h dark photoperiod throughout the experiment.

Fish were acclimated to 10°C, 15°C, or 20°C for at least one month before measuring their metabolic rate. Different individuals were used at each of these temperatures, and we used multiple tanks for each acclimation temperature. The intermediate temperature (15°C) is close to the maximum temperature experienced by fish in cold habitats in the summer and the minimum temperature experienced by fish in warm habitats in the winter (Table 1). The lowest temperature in this range (10°C) is not generally experienced by warm-habitat fish in the wild, and the highest temperature in this range (20°C) is not experienced by cold-habitat fish (Table 1). Exposing fish to these unfamiliar temperatures allowed us to examine the release of cryptic genetic variation in metabolic rate.

### Metabolic rate measurements

We used intermittent flow-through respirometry to estimate individual metabolic rates by measuring oxygen uptake. Sixteen cylindrical, borosilicate glass respirometry chambers (inner diameter: 32.3 mm; length: 124 mm; volume: 83 mL) were submerged in a 93-L experimental tank (780 mm × 570 mm × 210 mm) containing air-saturated water. The water temperature within the experimental tank was maintained at 10°C, 15°C, or 20°C depending on the treatment. This was done using a thermostated reservoir connected to the experimental tank by a thermoregulator (TMP-REG system, Loligo Systems, Denmark), which allowed us to maintain the water temperature within 0.2°C of our target temperature for the entire 24-h trial period. To maintain good water mixing and avoid an oxygen gradient in the respirometry chambers, we used a peristaltic pump (Masterflex, Cole-Parmer), which moved water through the chambers and around an external circuit of gas-impermeable tubing (Masterflex, Cole-Parmer). Oxygen concentration in the chambers was measured every two seconds using four Firesting channel oxygen meters with sixteen associated sensors (PyroScience GmbH, Aachen, Germany). To account for bacterial respiration during the trials, background bacterial oxygen consumption was measured before and after each trial in the 16 respirometry chambers (see *Data analysis*). A UV filter connected to the experimental tank was also used to sterilise the water and minimise bacterial respiration.

We fasted fish for 48 hr before the trials because metabolic rate increases during digestion (Killen 2014). Trials started around 14:00 each day. Immediately before being placed into a respirometry chamber, each fish was subjected to exhaustive exercise by being chased in a circular tank; this allowed us to measure their maximum metabolic capacity (Killen et al. 2012, Clark et al. 2013, Killen et al. 2017). After complete exhaustion, which always occurred within 2–3 min of chasing, fish were placed into individual respirometers. Rates of oxygen uptake were then measured in 3-min intervals over a 15-min period, during which the respirometers were sealed and the decrease in oxygen content was used to calculate rate of oxygen uptake (see *Data analysis*). The maximum rate of oxygen uptake measured during these five 3-min intervals was used as a proxy for the maximum metabolic rate (MMR).

The fish were left in the respirometers undisturbed until around 14:00 the following day. Every 9 min, an automated water pump (Eheim GmbH & Co. KG, Germany) would switch on for 2 min flushing the respirometers with aerated water. Based on the decrease in oxygen concentration during the 7-min off-cycle of the pumps, we calculated the rate of oxygen uptake. Standard metabolic rate (SMR) was estimated as the lowest 10th percentile of measurements taken throughout the measurement period (Dupont-Prinet et al. 2010, Killen 2014), excluding the first 5 hr during which oxygen consumption was elevated due to handling stress (Killen 2014). Absolute aerobic scope (AAS) was calculated as the difference between SMR and MMR. Factorial aerobic scope (FAS) was calculated as the ratio of MMR and SMR.

During the trial period, the experimental bath was covered with black plastic to avoid external disturbances. We also covered the sides of the respirometry chambers with opaque material to prevent visual stimuli from other individuals in the same trial. Fish were removed from the respirometer after 24 hr, at which point we weighed them and returned them to their initial holding tank. Our sample sizes ranged between 15 and 33 per population per temperature (Supplementary Tables 1 & 2). Due to equipment failure during a small proportion of our trials, we have slightly more measurements of MMR than SMR and AS (Supplementary Tables 1 & 2).

### Data analysis

Oxygen concentration data derived from Firesting software were analysed in LabChart 7 (ADInstruments Pty Ltd, Australia). To calculate oxygen consumption rates (*MO_2_*), we used the average slope of each 7-min off-cycle measurement period derived from the linear regressions between oxygen consumption over time (mg O_2_ h^−1^). All *MO_2_* data were corrected for the volume of the respirometry chamber and body mass of the fish. Using the bacterial respiration data collected for all chambers before and after trial measurements, we assumed a linear increase in bacterial oxygen consumption over time. The estimated bacterial background respiration at any given timepoint was then subtracted from the *MO_2_* data.

Statistical analyses were performed using R version 3.5.1 (R Core Team 2017), and figures were generated using the ggplot2 package (Wickham 2009). SMR, MMR, AAS, FAS, and body mass were log-transformed for use in models because of a non-linear relationship between metabolic rate and mass. For each response variable (SMR, MMR, AAS, and FAS), we used general linear models (GLM) with the following explanatory variables: thermal habitat (warm or cold), population pair (allopatric, sympatric 1, or sympatric 2), acclimation temperature (10°C, 15°C, or 20°C), and all possible interactions between these three factors. A significant effect of the three-way interaction between thermal habitat, population pair, and acclimation temperature would indicate non-parallel divergence of metabolic rate reaction norms across population pairs (Bolnick et al. 2018). After checking for homogeneity of slopes, body mass was also included as a covariate in all models to correct for the effects of mass on metabolic rate. All model assumptions were tested and verified, and the statistical results reported below are the values from the full models including all interactions. Our results for AAS (Table 2, Figure 2C) versus FAS (Supplementary Table 3, Supplementary Figure 1) are largely similar, and we only present the results for AAS, which is considered to be more informative and robust (Halsey et al. 2018).

**Table 2.** Results of general linear models testing the effects of thermal habitat (warm or cold), population pair (allopatric, sympatric 1, or sympatric 2), acclimation temperature (10°C, 15°C, or 20°C), and their interactions on SMR, MMR, and AS in threespine stickleback from six populations in Iceland. Df denotes degrees of freedom. Eta-squared (η^2^) represents the percent variance explained by each factor, which was calculated by dividing the sum of squares for each factor by the total sum of squares and multiplying by 100.

|  | Standard metabolic rate (SMR) |  |  |  | Maximum metabolic rate (MMR) |  |  |  | Aerobic scope (AS) |  |  |  |
| --- | --- | --- | --- | --- | --- | --- | --- | --- | --- | --- | --- | --- |
| | $\eta^2$ | df | *F* | *P* | $\eta^2$ | Df | *F* | *P* | $\eta^2$ | df | *F* | *P* |
| Thermal habitat | 0.82 | 1 | 9.77 | **0.002** | 1.38 | 1 | 10.4 | **0.001** | 0.99 | 1 | 6.26 | **0.013** |
| Population pair | 1.38 | 2 | 8.25 | **<0.001** | 0.81 | 2 | 3.27 | **0.039** | 2.20 | 2 | 7.09 | **<0.001** |
| Acclimation temperature | 31.7 | 2 | 192 | **<0.001** | 6.92 | 2 | 26.9 | **<0.001** | 2.97 | 2 | 9.56 | **<0.001** |
| Mass | 29.3 | 1 | 355 | **<0.001** | 35.9 | 1 | 281 | **<0.001** | 28.6 | 1 | 186 | **<0.001** |
| Thermal habitat × population pair | 0.94 | 2 | 5.65 | **0.004** | 0.35 | 2 | 1.56 | 0.21 | 0.22 | 2 | 0.89 | 0.41 |
| Thermal habitat × acclimation temperature | 0.03 | 2 | 0.14 | 0.87 | 1.15 | 2 | 4.29 | **0.014** | 1.43 | 2 | 4.74 | **0.009** |
| Population pair × acclimation temperature | 3.32 | 2 | 10.0 | **<0.001** | 0.46 | 2 | 0.98 | 0.42 | 1.21 | 2 | 2.05 | 0.087 |
| Thermal habitat × population pair × acclimation temperature | 0.82 | 4 | 2.49 | **0.043** | 1.15 | 4 | 2.25 | 0.063 | 2.97 | 4 | 4.79 | **<0.001** |
| Error | 31.7 | 385 |  |  | 51.9 | 406 |  |  | 59.4 | 385 |  |  |

**Figure 2.**
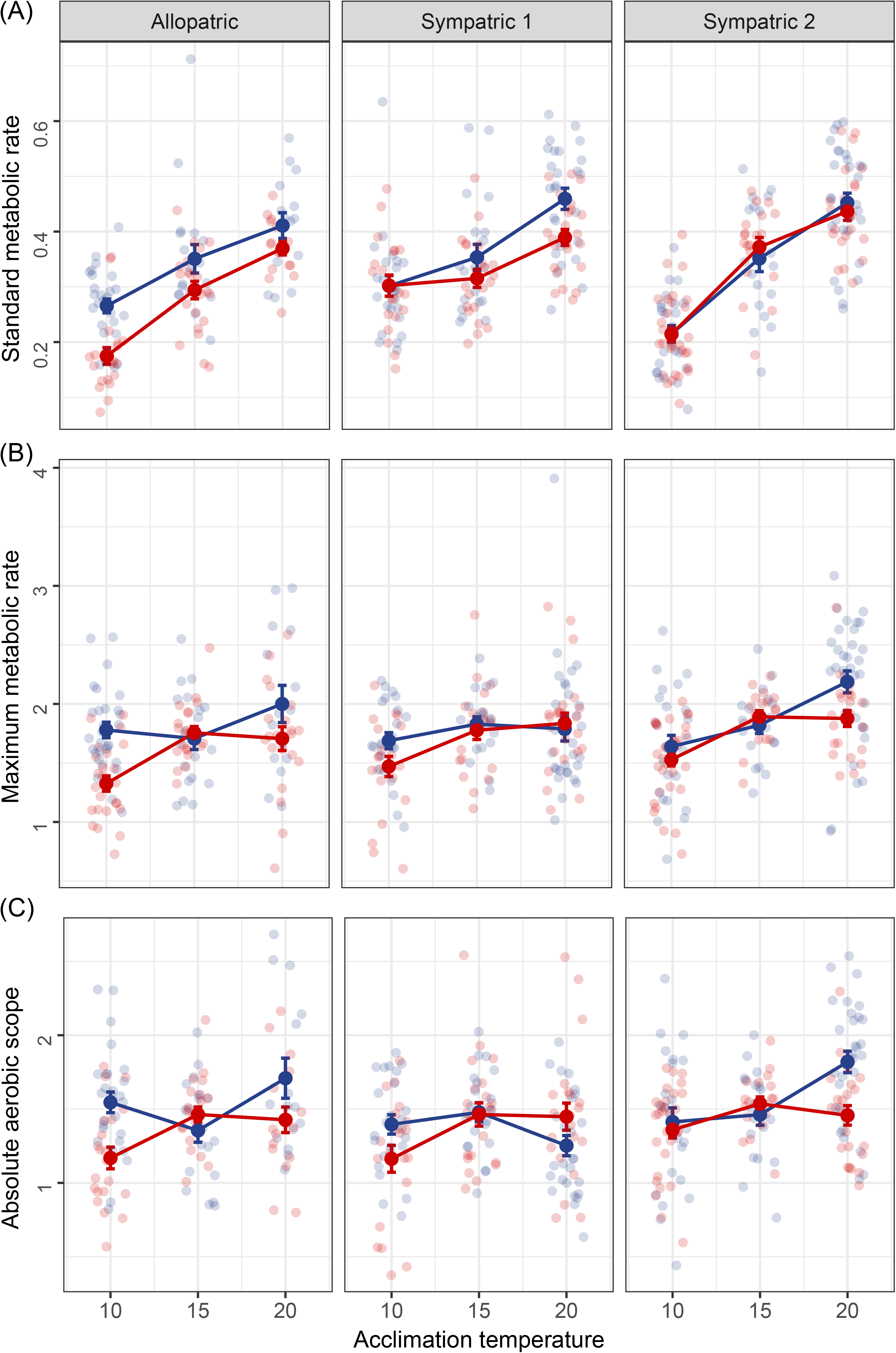
Metabolic rate (mg O_2_ hr^−1^) of threespine sticklebacks from cold and warm habitats in Iceland that were acclimated to 10°C, 15°C, or 20°C. We show mass-corrected standard metabolic rate (A), maximum metabolic rate (B), and aerobic scope (C). Error bars indicate standard errors, and small circles represent individual data points (blue=cold thermal habitat, red=warm thermal habitat). ‘Allopatric’ refers to Grettislaug and Garðsvatn, ‘sympatric 1’ refers to Áshildarholtsvatn, and ‘sympatric 2’ refers to Mývatn.

## Results

As expected, larger fish had higher absolute metabolic rates, and metabolic rate increased with acclimation temperature (Table 2). Acclimation temperature explained over 30% of the variation in SMR but only 7% and 3% of the variation in MMR and AAS, respectively (Table 2). SMR was therefore more variable than MMR and AAS in response to acclimation temperature. Fish from warm and cold habitats also differed in the steepness of their metabolic rate reaction norms (Figure 2). In terms of SMR, warm-habitat fish had a steeper metabolic rate reaction norm than cold-habitat fish in the allopatric population pair but a less steep reaction norm in sympatric population 1 (Figure 2A).

We found a statistically significant three-way interaction between thermal habitat, population pair, and acclimation temperature on SMR and AAS (Table 2). This interaction indicates that the divergence in metabolic rate reaction norms between warm- and cold-habitat fish varied across the three population pairs (Figure 2). For example, there was a stronger divergence in SMR between the warm- and cold-habitat fish in the allopatric populations than in the sympatric populations (Figure 2A). Similarly, the strong effect of population pair and acclimation temperature × population pair on SMR (Table 2) indicates that local adaptation may be driving variation in this trait across different sites.

Differences in SMR between fish from warm and cold habitats tended to be more pronounced at more extreme acclimation temperatures: in the allopatric population pair, thermal divergence in SMR decreased with increasing acclimation temperature, and in sympatric population 1, thermal divergence in SMR increased with increasing acclimation temperature (Figure 2A). However, in sympatric population 2, there was no divergence in SMR between warm- and cold-habitat fish at any of the three acclimation temperatures (Figure 2A).

Similar to SMR, differences in MMR between thermal habitats tended to be more pronounced at more extreme acclimation temperatures. Warm-habitat fish from sympatric population 2 had a lower MMR and AAS than cold-habitat fish when both were acclimated to 20°C (Figure 2B and 2C). In the other two population pairs, warm-habitat fish had a lower MMR and AAS than cold-habitat fish at 10°C (Figures 2B and 2C).

## Discussion

We have taken advantage of a unique study system of geothermally heated and ambient-temperature populations to test whether fish in warm environments show a suppressed or elevated metabolic rate compared to those in cold environments. We found a general pattern for a lower SMR in sticklebacks originating from warm habitats, although the extent of this effect varied depending on the population pair and acclimation temperature. Interestingly, the SMR of warm-habitat sticklebacks at their naturally experienced temperatures (15-20°C) was similar to the SMR of cold-habitat sticklebacks at their naturally experienced temperature (10-15°C). This results in similar metabolic costs in these contrasting thermal environments and is consistent with the predictions of countergradient variation and Krogh’s rule. Our findings therefore suggest that fish may evolve a lower metabolic rate as global temperatures increase in response to climate change.

In our study system, some populations of sticklebacks living in warm or cold habitats are in separate water bodies (allopatry), while others are found in different parts of the same water body (sympatry). Given the potential for gene flow between sympatric but not allopatric population pairs, we might expect sympatric pairs to be less phenotypically divergent than allopatric pairs (Hendry and Taylor 2004, Pinho and Hey 2010). Alternatively, sympatric pairs might be more divergent because of character displacement, whereby differences between morphs are more pronounced in areas where they co-occur and minimised in areas where their distributions do not overlap (Brown and Wilson 1956, Losos 2011). Our findings show a stronger divergence in metabolic rate between sticklebacks from warm and cold habitats in allopatry than in sympatry. This suggests that gene flow may constrain physiological adaptation in natural populations where physical dispersal between thermal habitats is possible (Lenormand 2002, Hendry and Taylor 2004). It also indicates that metabolic responses to thermal habitat vary across populations, making it difficult to predict metabolic rate evolution in response to climate change.

The two sympatric populations differed in the extent of thermal divergence in their metabolic rate reaction norms: sympatric population 1 showed a greater degree of divergence than sympatric population 2. It is possible that this variation is related to differences in the age of these warm habitats (Table 1). In populations that have been diverging for longer, there is more scope for natural selection and genetic drift to introduce adaptive or stochastic phenotypic differences (Ord and Summers 2015). In our study system, we might thus expect the younger population pair (i.e., sympatric 1) to be less divergent than the older populations pair (i.e., sympatric 2). However, we found the opposite pattern, where the young sympatric population showed a greater degree of divergence in metabolism between fish from warm and cold habitats. In this young sympatric population, there was thermal divergence in both SMR and MMR reaction norms, whereas in the older sympatric population, there was no divergence in SMR and only a small divergence in MMR. One explanation could be that there is low gene flow in the younger sympatric population and high gene flow in the older sympatric population. We believe this is unlikely given that the warm and cold habitats of the younger population are only tens of meters apart, and in the older one they are a few kilometres apart (Table 1). An alternative explanation is that there are differences in food availability in the two sympatric populations (White and Kearney 2013). Low food availability in the warm habitat of sympatric population 1, along with elevated temperatures, may favour a lower SMR to allow organisms to resist starvation (Alton et al. 2017). Correspondingly, if there is high food availability in the warm habitat of sympatric population 2 (O’Gorman et al. 2016), selection due to temperature and food availability might be operating in opposite directions, resulting in a similar SMR in the cold and warm habitats (Figure 2A).

Another interesting finding was a higher degree of variability in SMR than MMR and AS in response to acclimation temperature. Across the three warm-cold population pairs, acclimation temperature explained over 30% of the variation in SMR but only 7% and 3% of the variation in MMR and AS, respectively. If rising ambient temperatures cause a greater increase in SMR relative to MMR, this will lead to a decrease in AS (Donelson et al. 2011). Our results support this prediction, as warm-habitat sticklebacks tended to have a lower AS than cold-habitat sticklebacks at a high temperature (20°C). It has been proposed that the capacity to meet increased oxygen demands at elevated temperatures may determine the persistence of fish populations in a warming climate (Donelson et al. 2011; but see Lefevre 2016, Jutfelt et al. 2018), so it is important to better understand and predict changes in AS in response to temperature changes (Sinclair et al. 2016).

By measuring metabolic rates across a range of temperatures, rather than a single temperature as is typically done (Bruneaux et al. 2014), we were able to compare the steepness of the metabolic reaction norms of warm versus cold populations. In terms of SMR, warm-habitat sticklebacks had a steeper metabolic rate reaction norm than cold-habitat sticklebacks in the allopatric population pair but a less steep reaction norm in sympatric population 1. Despite these contrasting trends, a common pattern emerging from our results was that differences between warm- and cold-habitat sticklebacks tended to be more pronounced at more extreme temperatures (10 °C or 20°C). This suggests that cryptic genetic variation was released upon exposure to these novel conditions, thus revealing hidden evolutionary potential (Paaby & Rockman 2014, Shama 2017). If we had only measured metabolic rates at an intermediate temperature, we would have detected little or no differences between warm- and cold-habitat sticklebacks. Our findings demonstrate that when studying individuals originating from different thermal environments in a common garden setting, it is much more informative to compare their reaction norms rather than use a single acclimation temperature (Bruneaux et al. 2014).

Moreover, using populations exposed to contrasting thermal habitats for many generations allowed us to examine long-term responses to elevated temperatures (Huss et al. 2019). This avoids the limitations of short-term laboratory experiments that expose animals to high temperature for only one or a few generations. Nevertheless, using wild populations presents other limitations, since factors other than the thermal habitat could be contributing to the differences we observed in the allopatric population pair. Data on additional allopatric populations are thus needed to determine whether metabolic rate evolution in response to temperature is repeatable.

We also note that the relative contribution of genetic change and plasticity in driving metabolic rate differences in this study system is still unknown. A recent review suggested that metabolic rate is generally highly heritable, although active metabolic rate tends to be more heritable than resting metabolic rate (Pettersen et al. 2018). Further work on the heritability of metabolic traits in this system would allow us to assess their potential to respond to selection.

In summary, we have shown a general pattern for a lower metabolic rate in fish from warm habitats, which provides a powerful test of and support for the controversial Krogh’s rule. We also found evidence for a stronger divergence in metabolic rate between warm and cold habitats in allopatry than sympatry, suggesting that gene flow may constrain physiological adaptation when dispersal between thermal habitats is possible. By focusing on natural populations living in contrasting thermal environments over a small geographic scale, our study offers valuable insights into how fishes and other ectotherms might physiologically adapt to global climate change.

## Acknowledgements

We thank Joseph Humble for his assistance with animal husbandry. We also thank Iain Hill, Tiffany Armstrong, Anna Persson, and Kári Heiðar Árnason for their help with fieldwork in Iceland. We are also grateful to Craig White, Fredrik Jutfelt, and an anonymous reviewer for their helpful comments on this manuscript.

## Ethical statement

Our study adheres to the ASAB/ABS Guidelines for the Use of Animals in Research, the institutional guidelines at University of Glasgow, and the legal requirements of the UK Home Office (Project License P89482164).

## Authors’ Contributions

NP, SSK, NBM, JL, and KJP conceived and designed the study. NP performed the experiment, analysed the data, and drafted the manuscript. BKK and SS provided supplies and helped coordinate fieldwork. All authors edited the manuscript and gave final approval for publication.

