## Supplementary Figure 1 for "Multigenerational exposure to elevated temperatures leads to a reduction in standard metabolic rate in the wild"

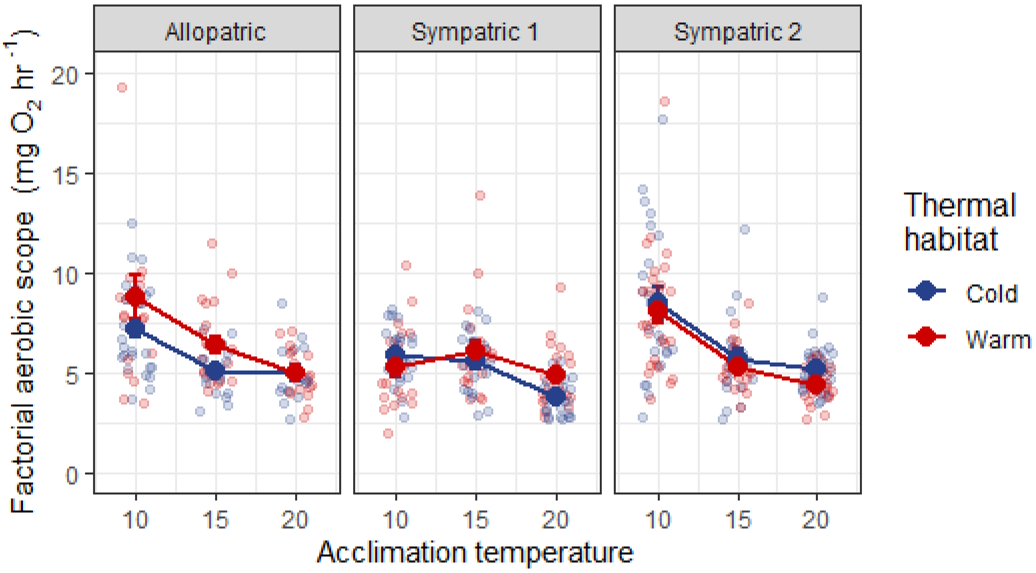


**Supplementary Figure 1.** Factorial aerobic scope (mg $O_2$ $hr^{-1}$) of threespine sticklebacks from cold and warm habitats in Iceland that were acclimated to 10°C, 15°C, or 20°C. Error bars indicate standard errors, and small circles represent individual data points (blue=cold thermal habitat, red=warm thermal habitat). 'Allopatric' refers to Grettislaug and Garðsvatn, 'sympatric 1' refers to Áshildarholtsvatn, and 'sympatric 2' refers to Mývatn.
