## Supplementary Table for "Multigenerational exposure to elevated temperatures leads to a reduction in standard metabolic rate in the wild"

**Supplementary Table 1.** Samples sizes for standard metabolic rate (SMR), absolute aerobic scope (AAS), and factorial aerobic scope (FAS) measurements at three acclimation temperatures (10 °C, 15 °C, and 20 °C). 'Allopatric' refers to Grettislaug and Garðsvatn, 'sympatric 1' refers to Áshildarholtsvatn, and 'sympatric 2' refers to Mývatn.

| Population pair | Thermal habitat | 10 °C | 15 °C | 20 °C |
|---|---|---|---|---|
| Allopatric population | Warm | 21 | 22 | 18 |
| | Cold | 27 | 19 | 15 |
| Sympatric population 1 | Warm | 20 | 21 | 23 |
| | Cold | 24 | 20 | 26 |
| Sympatric population 2 | Warm | 30 | 19 | 23 |
| | Cold | 24 | 19 | 31 |

**Supplementary Table 2.** Sample sizes for maximum metabolic rate (MMR) measurements at three acclimation temperatures (10 °C, 15 °C, and 20 °C). 'Allopatric' refers to Grettislaug and Garðsvatn, 'sympatric 1' refers to Áshildarholtsvatn, and 'sympatric 2' refers to Mývatn.

| Population pair | Thermal habitat | 10 °C | 15 °C | 20 °C |
|---|---|---|---|---|
| Allopatric population | Warm | 24 | 22 | 21 |
| | Cold | 29 | 19 | 15 |
| Sympatric population 1 | Warm | 22 | 22 | 24 |
| | Cold | 24 | 20 | 29 |
| Sympatric population 2 | Warm | 35 | 19 | 23 |
| | Cold | 25 | 19 | 33 |

**Supplementary Table 3.** Results of general linear model testing the effects of thermal habitat (warm or cold), population pair (allopatric, sympatric 1, or sympatric 2), acclimation temperature (10°C, 15°C, or 20°C), and their interactions on factorial aerobic scope (FAS) in threespine stickleback from six populations in Iceland. Df denotes degrees of freedom. Eta-squared ($\eta^2$) represents the percent variance explained by each factor, which was calculated by dividing the sum of squares for each factor by the total sum of squares and multiplying by 100.

| | Facultative aerobic scope (FAS) | | | |
|---|---|---|---|---|
| | $\eta^2$ | df | *F* | *P* |
| Thermal habitat | 0.09 | 1 | 0.51 | 0.47 |
| Population pair | 3.94 | 2 | 11.2 | **<0.001** |
| Acclimation temperature | 21.8 | 1 | 124 | **<0.001** |
| Mass | 1.80 | 1 | 10.2 | **<0.001** |
| Thermal habitat × Population pair | 0.86 | 2 | 2.43 | 0.089 |
| Thermal habitat × Acclimation temperature | 0.09 | 1 | 0.54 | 0.46 |
| Population pair × Acclimation temperature | 1.29 | 2 | 3.67 | **0.026** |
| Thermal habitat × Population pair × Acclimation temperature | 1.64 | 2 | 4.66 | **0.010** |
| Error | 68.5 | 385 | | |
